# Variable rates of SARS-CoV-2 evolution in chronic infections

**DOI:** 10.1101/2024.05.31.596818

**Authors:** Ewan Smith, William L. Hamilton, Ben Warne, Elena R. Walker, Aminu S. Jahun, Myra Hosmillo, ISARIC Consortium, Ravindra K. Gupta, Ian Goodfellow, Effrossyni Gkrania-Klotsas, M. Estée Török, Christopher J. R. Illingworth

**Affiliations:** MRC-University of Glasgow Centre for Virus Research, Glasgow, UK; Department of Medicine, University of Cambridge, Cambridge, United Kingdom; Cambridge University Hospitals NHS Foundation Trust, Cambridge, United Kingdom; Wellcome Sanger Institute, Wellcome Trust Genome Campus, Hinxton, United Kingdom; Division of Virology, Department of Virology, University of Cambridge, Cambridge, United Kingdom; Cambridge Institute of Therapeutic Immunology & Infectious Disease (CITIID), Cambridge, UK; MRC Epidemiology Unit, University of Cambridge, Level 3 Institute of Metabolic Science, Cambridge, United Kingdom; School of Clinical Medicine, University of Cambridge, Cambridge, United Kingdom

## Abstract

An important feature of the evolution of the SARS-CoV-2 virus has been the emergence of highly mutated novel variants, which are characterised by the gain of multiple mutations relative to viruses circulating in the general global population. Cases of chronic viral infection have been suggested as an explanation for this phenomenon, whereby an extended period of infection, with an increased rate of evolution, creates viruses with substantial genetic novelty. However, measuring a rate of evolution during chronic infection is made more difficult by the potential existence of compartmentalisation in the viral population, whereby the viruses in a host form distinct subpopulations. We here describe and apply a novel statistical method to study within-host virus evolution, identifying the minimum number of subpopulations required to explain sequence data observed from cases of chronic infection, and inferring rates for within-host viral evolution. Across nine cases of chronic SARS-CoV-2 infection in hospitalised patients we find that non-trivial population structure is relatively common, with four cases showing evidence of more than one viral population evolving independently within the host. We find cases of within-host evolution proceeding significantly faster, and significantly slower, than that of the global SARS-CoV-2 population, and of cases in which viral subpopulations in the same host have statistically distinguishable rates of evolution. Non-trivial population structure was associated with high rates of within-host evolution that were systematically underestimated by a more standard inference method.

## Introduction

The evolution of the SARS-CoV-2 virus during the early phases of the pandemic has been characterised by two apparently distinct processes^1^. In the first process, the viral population gained in diversity via the gradual accumulation of mutations. Mutations acquired in this process included those which conferred a fitness advantage upon the virus. For example, the D614G mutation in Spike protein was associated with higher viral load, and was observed at an increasing frequency over the course of 2020 within the UK^2^. The accumulation of mutations provided a distinct genetic signal, with the genetic distance of a virus from the original Wuhan strain increasing roughly linearly with time. In the second process, viral genomes were sometimes observed that differed by multiple mutations from anything else observed in the viral population. Notable examples of this were the Alpha and Omicron variants of concern^3,4^. These two processes have been understood as the combined result of within-host and between-host virus evolution. For example a viral population acquiring genetic substitutions during the course of a chronic case of infection might remain unobserved for a substantial period of time before spilling into the general population via a transmission event^5,6^. Multiple individual cases of chronic SARS-CoV-2 infection have been described in the literature^7–9^. In one example, the Δ69-70 deletion in the Spike protein, characteristic of the Alpha variant, was observed prior to the emergence of that variant during a case of chronic infection^10^.

The hypothesis that chronic infection with SARS-CoV-2 lay behind the emergence of both the Alpha and Omicron strains imposes two demands upon cases of chronic infection. Firstly, to produce highly mutated viruses, evolution must proceed faster during the period of chronic infection than it would in a pattern of regular cases of infection and transmission, generating and fixing multiple substitutions in the viral genome. Secondly, having acquired these mutations, the virus must transmit back into the regular population, maintaining sufficient fitness to compete with viruses that had not passed through a chronically infected host. Positive selection, either for immune escape or to evade antiviral therapy, provides one mechanism via which rapid within-host evolution could occur. A study of multiple cases of SARS-CoV-2 evolution in immunocompromised hosts identified an increased proportion of substitutions in the Spike protein compared to the global viral population, supporting the hypothesis of positive selection^11^. Other analyses have identified increased burdens of mutation in viral populations from immunocompromised hosts^12,13^, incidences of transmission of SARS-CoV-2 from cases of chronic infection^14^, the emergence in immunocompromised hosts of variants conferring immune evasion^15^, and the early emergence of variants yet to be found in the global viral population^16^.

Although the existence of positive selection during SARS-CoV-2 infection is clear, the presence of selection does not imply an increased rate of virus evolution. Sequence data from selected cases of chronic SARS-CoV-2 infection have been used to identify more rapid rates of evolution during within-host infection than in the global population^8,17^. However, the overall picture is complex. An analysis of more than 100 cases of SARS-CoV-2 infection in hospitals did not find a generally increased rate of within-host evolution^18^. A recent overview of several hundred cases of chronic infection found examples of both apparently low and apparently high rates of within-host evolution^19^.

Estimating within-host rates of virus evolution is a complex task, with multiple factors potentially confounding the calculation. Methods for phylogenetic analysis, used for evolutionary rate estimation in global RNA viral populations^20^, assume that the sequences which make up the tree describe the genomes of individuals in a population^21^. When applied to a set of consensus sequences from a within-host viral population, this assumption is not correct. Phylogenetic methods usually do not account for errors in sequencing, which can increase terminal branch lengths^22^. Perhaps most significantly, efforts to infer within-host evolutionary rates rarely include a formal consideration of within-host population structure. Population structure has been hypothesised as contributing factor to the persistence of SARS-CoV-2 infection^23^, with subpopulations potentially existing deep within the lungs^24^ or elsewhere in the body^25^. For the purposes of rate estimation this complicates matters, in so far as distinct within-host populations may evolve at different rates^26^. A single rate of evolution, however inferred, may not reflect biological reality.

Here we use two novel approaches to gain an insight into within-host SARS-CoV-2 evolution. A first approach conducts sequence-based cartography, mapping consensus sequences, and therefore patterns of within-host evolution, in a simple visual manner. While only semi-quantitative in nature, this approach highlights potentially complex patterns of virus evolution, including potential substructure in the population, in cases of virus infection. A second approach performs a formal statistical calculation to infer the number of distinct viral populations within a host from what are assumed to be error-prone data, estimating rates of evolution for each population identified. Examining a set of 9 cases of chronic SARS-CoV-2 infection, we find evidence that within-host population structure, involving independently-evolving viral populations, is a common feature of infection. Subpopulations within the same host are found to evolve at significantly distinct rates, with a complex relationship pertaining between selection, viral phenotype, and the within-host rate of evolution.

## Results

Genome sequence data described cases of chronic infection in nine hospitalised individuals. Cases were selected under the criteria of having had viral genome sequences collected on at least four independent occasions, with these sequences spanning a period of infection of at least 21 days. Genome sequence data from one case (patient H) were described in an earlier publication^10^. The remaining cases originated from early in the pandemic, between March and June 2020. In this period, vaccination against SARS-CoV-2 had not been introduced^27^, while specific antiviral treatments were few.

An initial assessment of sequence data from these cases provided a qualitative insight into the structures of the different viral populations (Figure 1). Phylogenies and sequence maps highlighted different patterns of evolution in different hosts. For example, in patient A, no change in the viral consensus sequence was observed over 39 days of observation. In patient B only a single change in the sequence was observed, with the consensus from the first sample differing by a single nucleotide from the remaining sequences; one sequence contained an ambiguous nucleotide at this variant position. Patients G and H showed complex patterns of evolution, with multiple branches diverging from the sequence collected in the first sample, potentially representing independent subpopulations Sequence maps showed information additional to the phylogenies, representing genome sequences in two-dimensional space. These plots capture uncertainties encoded in ambiguous nucleotides, and highlighted pattens of divergent evolution. Phylogenies and sequence maps for other individuals are shown in Supplementary Figure 1.

**Figure 1:**
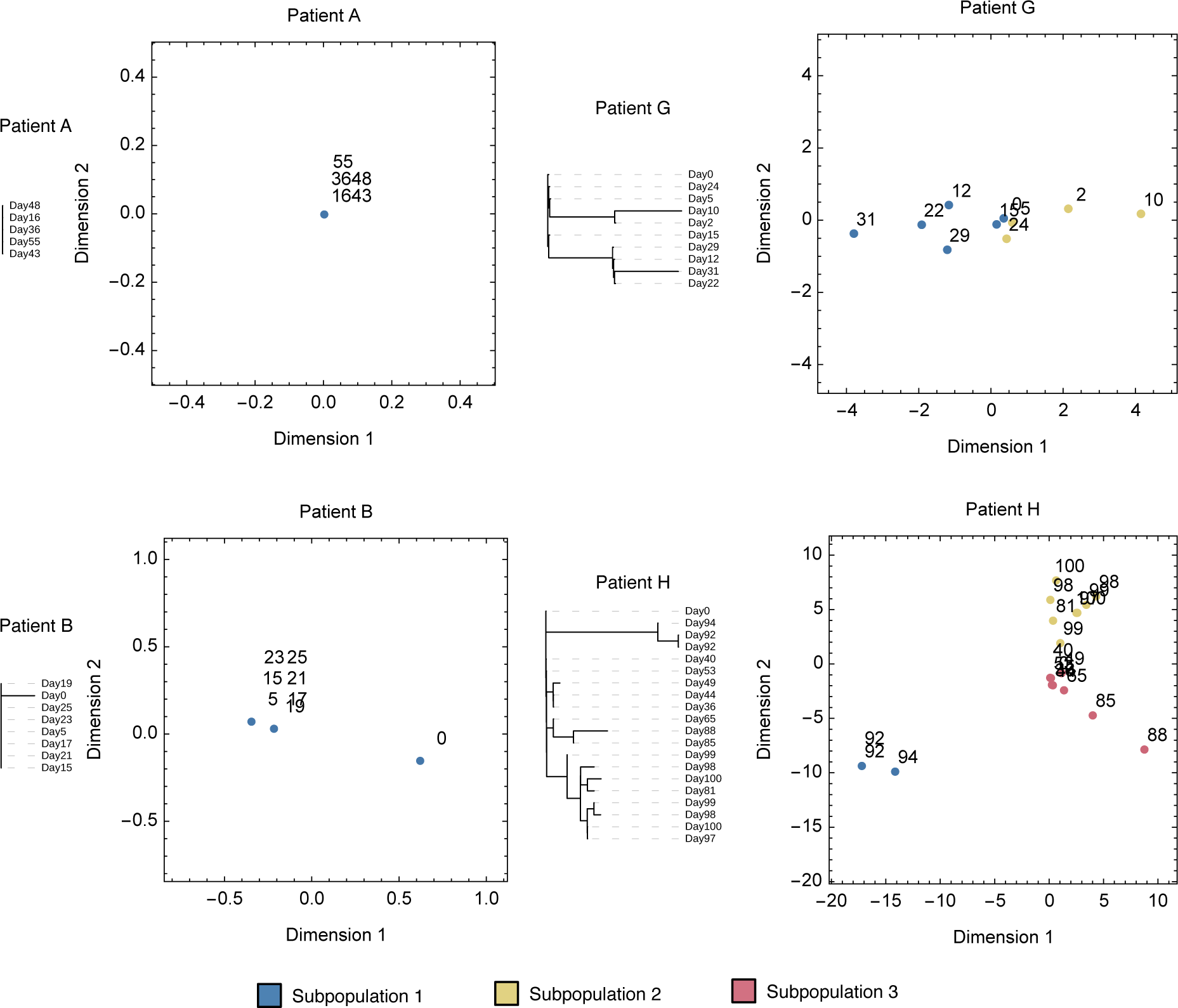
Phylogenetic plots and sequence maps for viral genome sequences collected from four patients in our study. In the sequence maps genome sequences are plotted as points in two dimensions, with numbers adjacent to points indicating the day on which corresponding samples were collected. Points are coloured according to the viral subpopulation they were assigned to by our inference method. The locations of points were calculated so that the pairwise spatial distances between points most closely approximate the pairwise Hamming distances between sequences, with a single unit in space corresponding to one nucleotide difference between sequences. Sequence maps explicitly incorporate information from ambiguous nucleotides at variant sites, leading to the potential for points to be separated by distances of less than one unit.

Where trees and sequence maps provided a qualitative insight into the relationships between samples from each host, a novel approach to the sequence data provided a formal insight into within-host population structure. Our approach fits models of different numbers of underlying populations to data from a host, identifying the smallest number of viral independent subpopulations required to explain the sequence data. Applied to simulated data our method showed a consistent ability to infer rates of viral evolution (Supplementary Figures 2 to 5), while being conservative in the number of subpopulations identified. While for a given host, distinct subpopulations were not always identified, overestimates of the number of subpopulations were rare, with one overestimate in 700 simulated populations.

Applied to our patient data, we inferred the presence of non-trivial population structure in four out of nine cases. In three of these cases two subpopulations were inferred to exist, while in one case three subpopulations were inferred. In patients G and H viral material was collected using different sample types (e.g. swab or sputum sample). No significant relationships were identified between sample type and the assignment of samples to subpopulations (Supplementary Tables 1 to 3). Data from the remaining cases was consistent with a single underlying viral subpopulation.

Rates of evolution estimated by our approach showed a high degree of variability (Figure 2). Compared to an estimated rate of evolution of 6 × 10^-4^ substitutions per site per year for hospitalise SARS-CoV-2 patients^18^, and an estimated rate of 8 × 10^-4^ substitutions per site per year for the global viral population^28^, maximum likelihood rates from our model lay between zero and 3.9 × 10^-3^ substitutions per site per year. In patients C, F, and H we identified viral populations evolving significantly faster than the global rate, with a 95% confidence interval in the estimated rate for a subpopulation excluding the global rate of evolution. Conversely in patient A we identified a single population evolving significantly slower than the global rate. Where more than one subpopulation was identified in a host, estimated rates of evolution for the faster-evolving population were between 1.9 and 4.9 times that of the global population. In two of the four cases where non-trivial population structure was identified evidence was found for different subpopulations having significantly different rates of evolution, suggesting that the virus evolved at different rates within the same host (Figure 2B).

**Figure 2:**
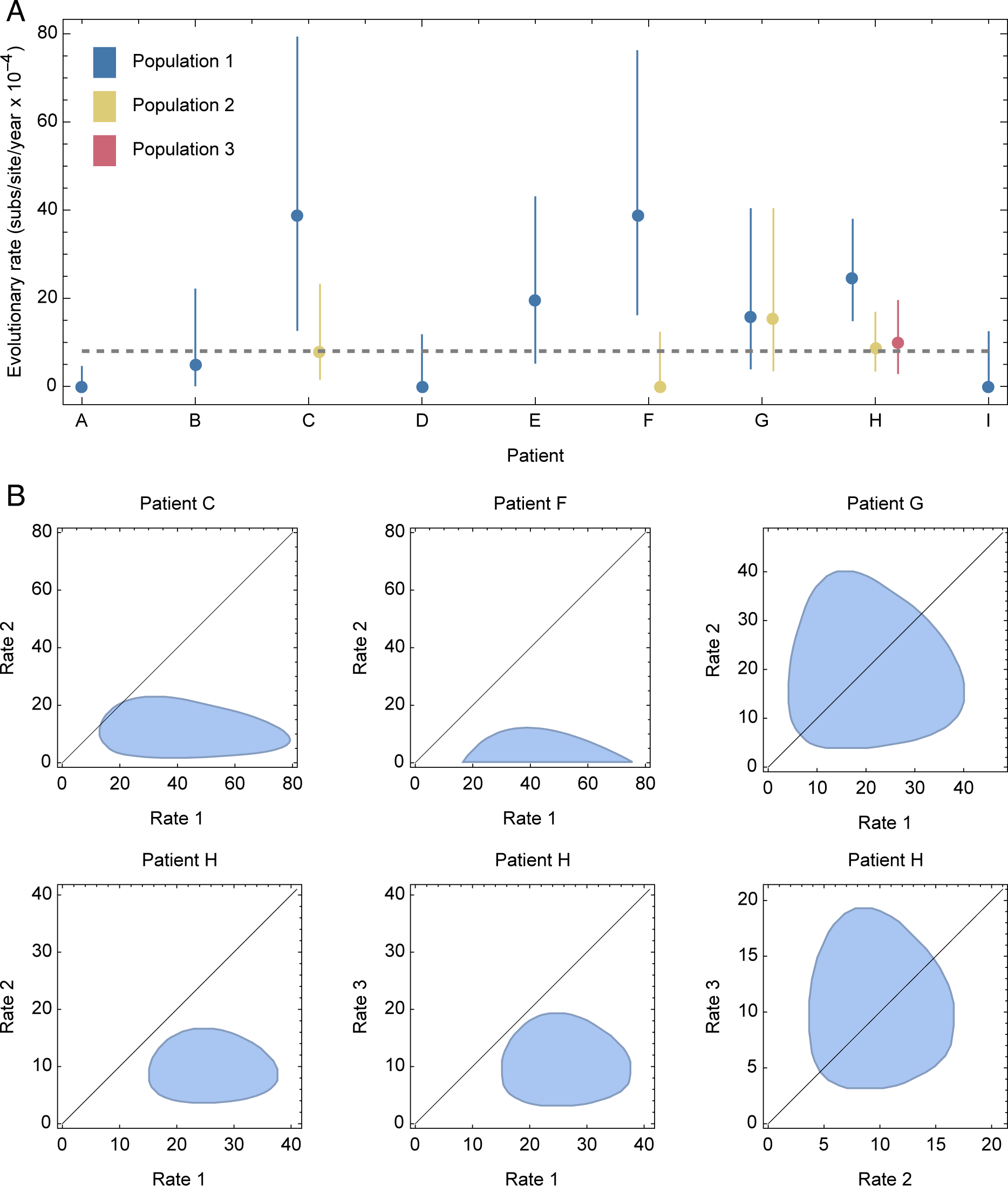
Inferred rates of within-host SARS-CoV-2 evolution. **A.** Maximum likelihood rates of within-host evolution are shown as dots, coloured according to the population identified within an individual. Between 1 and 3 populations were identified per host from the data. Vertical lines show estimated uncertainties in each rate. The horizontal gray dashed line shows an estimate for the global rate of SARS-CoV-2 evolution. **B.** Uncertainties in joint estimates of rates for individuals in which more than one population was identified. The black line shows parity between rates of evolution.

We further estimated rates of evolutionary change at synonymous and non-synonymous sites in the genome. Inferred rates of evolution at synonymous sites were generally faster than those at non-synonymous sites (Figure 3A). Among populations for which both synonymous and non-synonymous substitutions were observed, the combined rate of evolution at these sites (i.e. excluding changes at non-coding positions) was strongly correlated with the rate of gain of non-synonymous mutations (Figure 3B). However, within these populations there was no significant link between a faster rate of evolution and a higher value of the statistic dN/dS (Figure 3C), as might arise if increased selection for non-synonymous change increased the rate of evolution. One challenge in measuring selection in this context is the small number of variants that arose in each individual; our individual-level dN/dS statistics may not be supported by enough data to identify underlying relationships between variant characteristics and evolutionary rate. However, in our dataset, populations in which greater proportions of non-synonymous mutations were observed were not associated with faster patterns of viral evolution.

**Figure 3:**
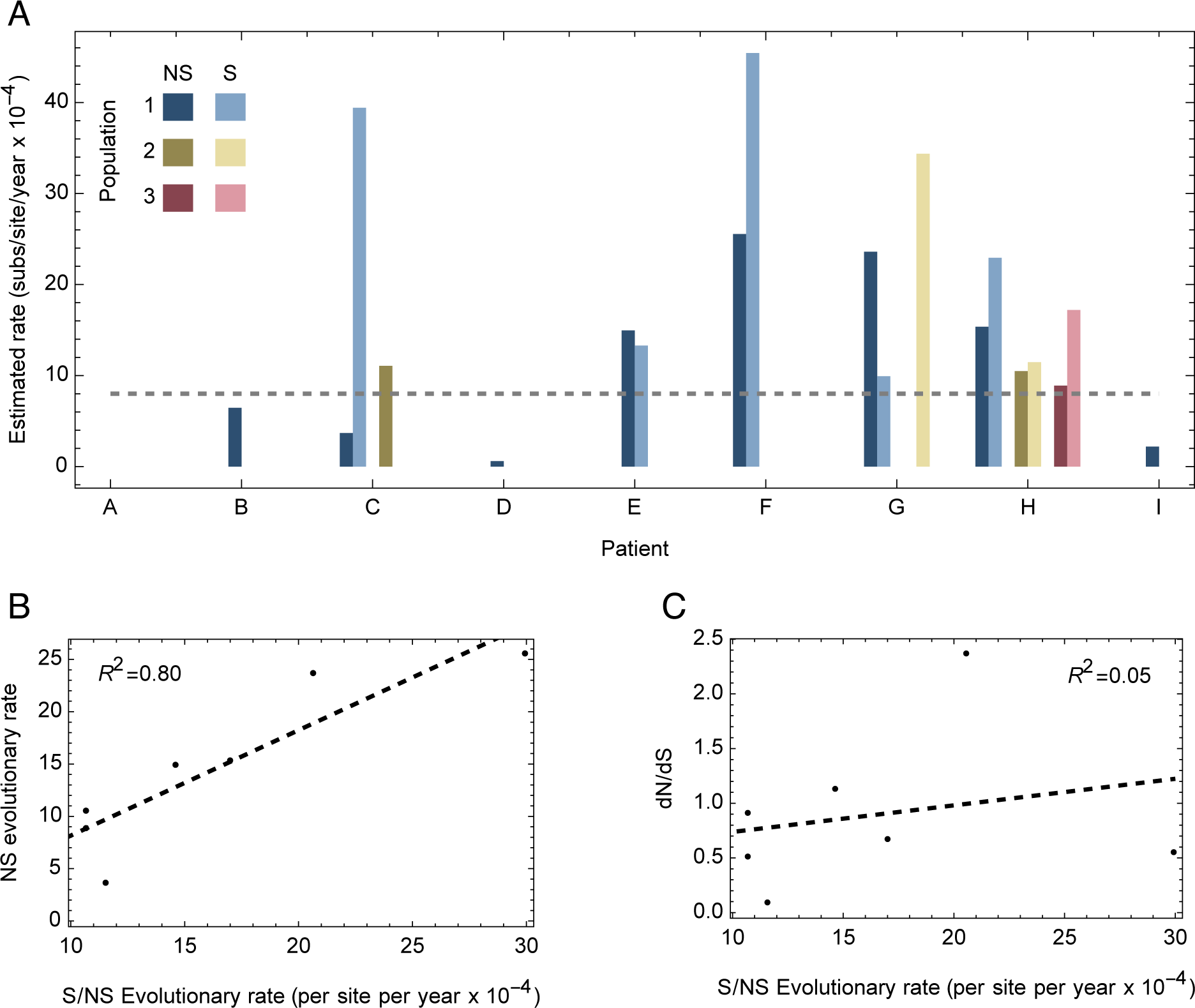
Inferred rates of evolution at synonymous and non-synonymous sites. **A.** Rates of evolution at synonymous and nonsynonymous sites were calculated across a statistical ensemble of model outputs. The horizontal dashed line shows an estimate of the global rate of SARS-CoV-2 evolution. **B.** Correlation between the inferred rate of evolution at nonsynonymous sites, and the total evolutionary rate calculated across both nonsynonymous and synonymous sites. The dashed black line shows a linear model fit to the data. **C.** Relationship between dN/dS and the total evolutionary rate calculated across both nonsynonymous and synonymous sites. The dashed black line shows a linear model fit to the data.

Our data were consistent with a higher rate of substitutions in the Spike protein than in the remainder of the genome^11,19^. Due to the design of our model, nucleotide substitutions could be inferred to occur with probabilities strictly between zero and one. We identified an expected 9.1 substitutions in Spike, with an expected 46.4 substitutions in the rest of the genome. Together these suggest a substitution rate for Spike that was 33% higher than for the rest of the genome (Figure 4), although the difference was not great enough to be statistically significant. Indeed, the locations of inferred sites of nucleotide substitutions in the genome were consistent with a uniform distribution across the genome (p=0.328, Kolmogorov Smirnov test). Roughly even distributions of variants have been noted for substitutions in the global SARS-CoV-2 population^11^, and for SARS-CoV-2 in hamsters following treatment with mutagenic drugs^29^.

**Figure 4:**
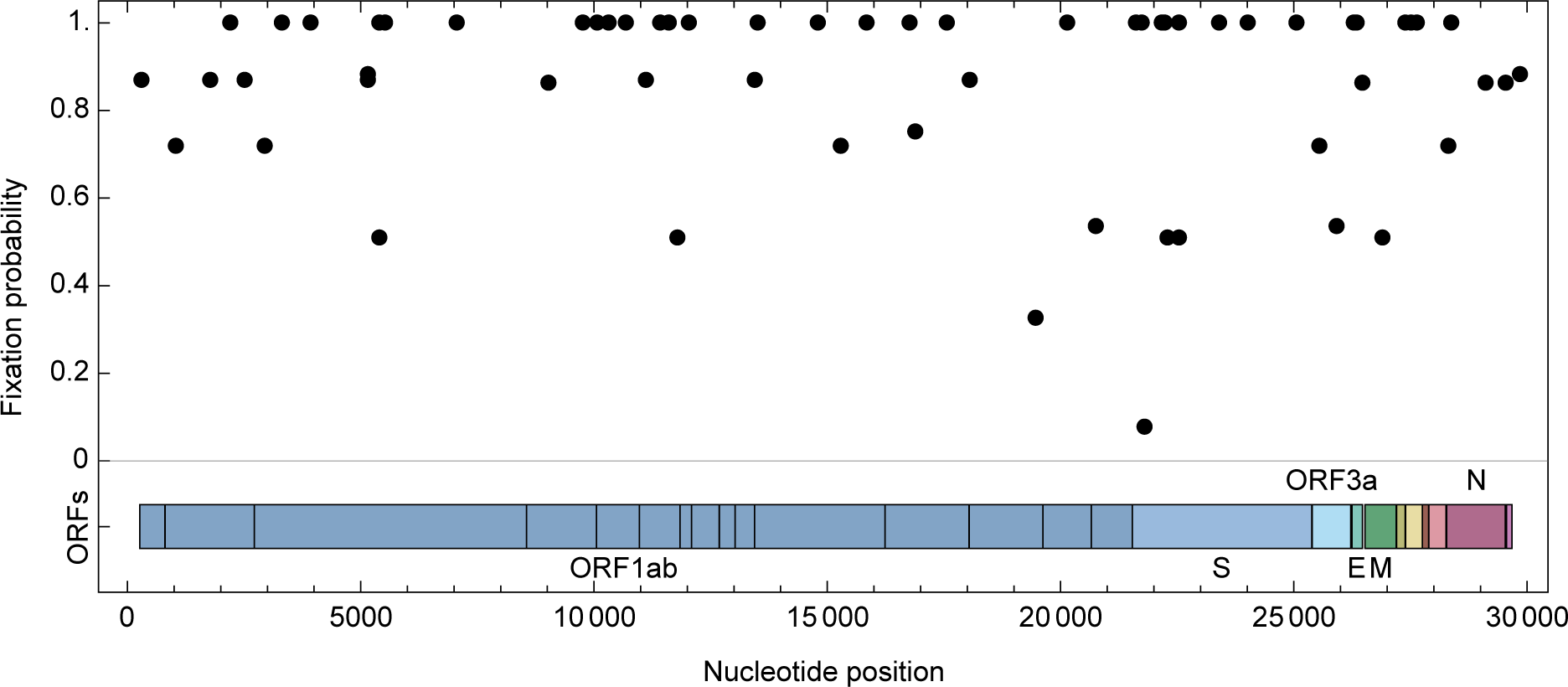
Locations in the genome of potential fixation events. Fixation events in our model are associated with a probability, which was calculated across an ensemble of models in which changes in the viral sequence could reflect either genuine change in a population or a form of sequencing error.

We combined the consensus sequences in our dataset with publicly-available short-read data to re-evaluate patterns of virus evolution in patient H. In the original analysis of this case, changes in allele frequencies showed rapid fluctuations in the frequencies of variants in the Spike protein. For example, viruses carrying the D796H mutation and a deletion of positions 69 and 70 (Δ69-70) rose to high frequency on two occasions shortly following the use of convalescent plasma therapy (Figure 5A). Our analysis of these data suggested that the observed fluctuations in allele frequencies could be explained in terms of competition between stable subpopulations, with each subpopulation gaining distinct fixations in the Spike protein over time (Figure 5B). Rather than observing dramatic changes in a well-mixed population, the evolutionary dynamics of the system reflect the dominance at different time of different subpopulations, either due to a selective response to variable antiviral therapy, or due to stochastic events in the emission and sampling of viral material from the host. We note that in this patient the D796H and Δ69-70 variants were associated with escape from convalescent plasma, while the Δ69-70 variant was one of the variants that defined the Alpha SARS-CoV-2 lineage which emerged in the global population some months after the clinical case^30^. Our decomposition identified both variants with a subpopulation in the host that was inferred to have a rate of evolution similar to that of the global population. In this case, the gain of variants of experimentally-verified phenotypic importance was not associated with an increased rate of viral evolution.

**Figure 5:**
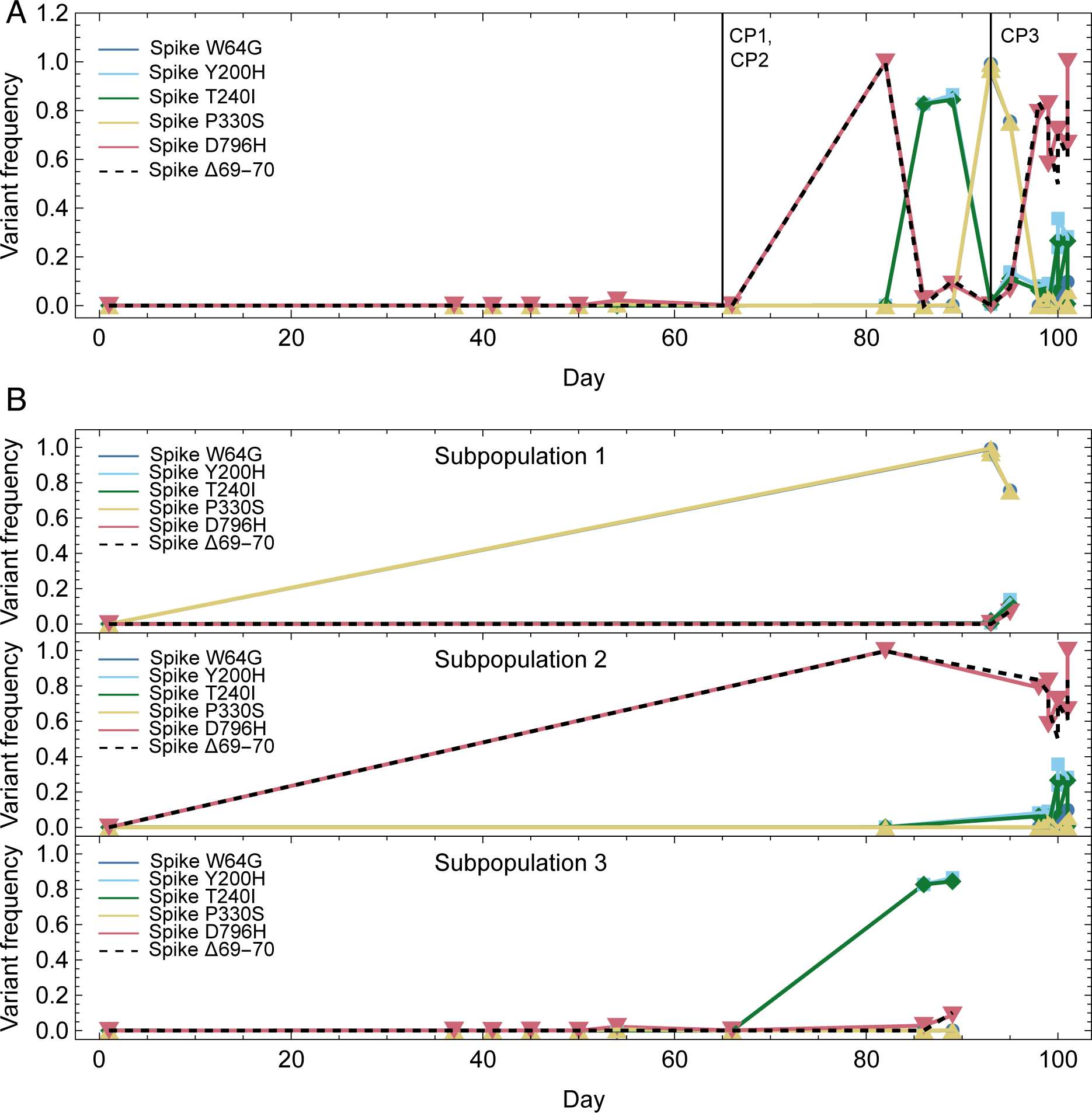
Changes in allele frequencies in the data from patient H. **A.** Allele frequencies of selected variants in the SARS-CoV-2 Spike protein, calculated from short-read sequence data from patient H. Vertical lines show the times of administration of three doses of convalescent plasma. **B.** Allele frequencies replotted according to a division of samples into subpopulations, as inferred by our method. Markers show the times of individual samples; lines connecting variant frequencies are for illustration only.

Our analysis of rates of evolution was repeated using a regression-based approach applied in other studies of within-host virus evolution^17,26^. This simpler approach, examining sequence divergence from the initial sample over time, could not capture the details of within-host population structure, for example inferring for patient F a rate intermediate to the rates of the different subpopulations we identified, or for patients C and G inferring a rate consistent with one of the subpopulations identified by our approach, but not consistent with another subpopulation in the same patient (Figure 6, Supplementary Figure 6). Of note, rates estimated with the simpler regression model were universally lower than that of the faster-evolving subpopulation in each host where more than one subpopulation was identified; previous studies may have underestimated rates of within-host evolution through not accounting for population structure.

**Figure 6:**
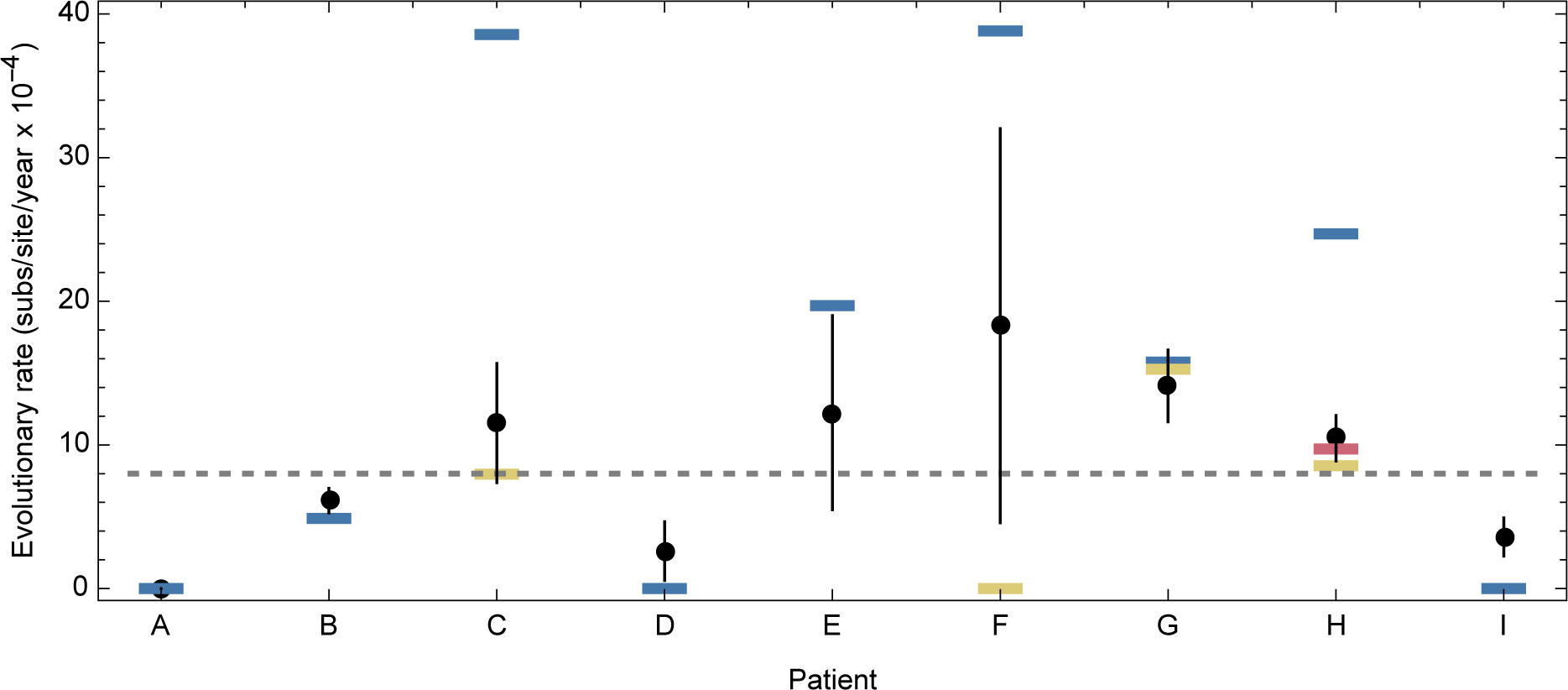
Rates of evolution inferred using a simple method of linear regression. The black dot for each patient shows an estimate of the within-host rate of virus evolution, inferred from a method of linear regression. Vertical black lines show error bars for these estimates. Blue, yellow, and red horizontal lines show the rates for the first, second, and third populations inferred by our approach. The linear regression method commonly underestimates the more rapid rates of evolution identified in structured cases of within-host evolution.

Clinical characteristics of the individuals in our study were explored but are not described in detail to preserve patient anonymity. Of the four patients in whom non-trivial population structure was identified, three were identified as being significantly immunocompromised, with conditions including primary immunodeficiency syndrome, haematological malignancy or organ transplantation. One of the remaining five patients was identified as being significantly immunocompromised. A previous study identified an increased rate of nucleotide substitutions in severely immunocompromised versus mildly or non-immunocompromised individuals^31^. Although in our study significantly immunocompromised individuals were more likely to have more complex population structure, the numbers involved were too small to achieve significance.

## Discussion

Within-host SARS-CoV-2 evolution may be more complex, and more varied, than has to date been appreciated. We have here applied a novel approach to study data from nine cases of chronic SARS-CoV-2 infection. Our method allows for a formal discrimination to be made between simple and complex population structures, inferring the presence of one or more independent viral subpopulations during a case of infection. Here, evidence for more than one subpopulation was found in four out of nine cases. We identified subpopulations evolving significantly faster, and significantly slower, than an estimated rate of evolution for the global SARS-CoV-2 population. Rates of evolution were not linked in a simple manner to viral phenotype. For example, phenotypically validated mutations identified in patient H and associated with escape from convalescent plasma therapy did not arise in a subpopulation with a higher inferred rate of evolution.

Our study is not the first to identify subpopulations during within-host SARS-CoV-2 evolution, though our method has novelty in doing this in a formal statistical manner. In identifying distinct subpopulations, our method provides an advantage over simple linear-regression-based models. Distance metrics have in the past been used to identify sequences that may be associated with faster-evolving subpopulations^10^. Against these approaches, our method can identify different populations even where their rates of evolution cannot be distinguished. Indirect evidence, such as the partial onset of drug resistance, or a limited extent of interaction between distinct viruses in a host, has also been used to highlight potential within-host structure. While indicative, such data does not constitute formal statistical evidence^32,33^. While direct sampling from different anatomical locations in an infected individual provides the best evidence of spatial separation^34,35^, there are practical limitations in obtaining such samples in routine clinical care.

One limitation of our work is the use of only respiratory samples for the diagnosis of population structure. Viral proteins and RNA have been identified in multiple other organs within individuals, creating the potential for sequence diversity going beyond what was accessible to our study^36^. Our statistical method makes a variety of simplifying assumptions which may also affect our results. Firstly, while distinct subpopulations in our model have independent rates of viral evolution, the rate of evolution for a single population is held to be constant. As such, time-dependent changes in the rate of evolution, as may occur when antiviral therapy is used in treatment, cannot be inferred. In this respect, the rates of evolution reported by our approach represent mean rates averaged across the period of infection. Time-dependent selection has been identified in genomic data describing respiratory infection^37^, but the limited nature of the data in this case mandate a cautious approach towards more complex evolutionary models. Calculating mean rates of evolution potentially obscures mixtures of higher and lower rates of evolution than are shown; in this respect the high rates of evolution we report are conservative. Secondly, our model assumes that subpopulations are founded at the time of infection by a virus or viruses which share a common sequence. This approach reflects limited within-host sequence diversity during SARS-CoV-2 infection and a tight population bottleneck at the time of infection^38^. In the absence of higher numbers of samples it is difficult to distinguish subpopulations initiated by genetically distinct viruses from independently evolving populations founded by identical viruses. In so far as our data show variants accumulating in populations over time they support the hypothesis of virus evolution, as opposed to simple and persistent genetic diversity that existed from the foundation of each infection. Thirdly, our approach models errors in consensus sequences as occurring via a regular Poisson process. In this sense we do not explicitly account for cases in which individual sequences include large numbers of sequence errors. A potential example of this in our dataset is Patient F, in which all but one of the sequences collected are identical, with the distinct sequence containing seven changes in the consensus sequence; alternative models of sequencing error could change our interpretation of these data. Strictly speaking the concept of error in our model is a broad one, encompassing for example the temporary changes in consensus that might result from clonal competition^39,40^. A prior evaluation of high quality SARS-CoV-2 genome sequences did not show a clear link between CT score and the extent of sequencing error^18^. Fourthly, we assumed that the detection of virus in quantities sufficient to produce sequence data implies the existence of a replicating viral population. If this assumption were not correct, the presence of residual viral genetic material could potentially explain apparently slower rates of evolution in some cases. We note that, if later samples from individuals are merely emissions from a dead virus population, the rates of evolution we report underestimate those for replicating virus populations. Finally, we assumed that each consensus sequence could be assigned uniquely to a single subpopulation. This assumption is supported by past evidence from cases of population structure in respiratory virus infection, which have suggested an either-or mechanism of emission, whereby each sample represents a discrete subpopulation, rather than a mixture of what is contained in a host^10,26^. However, not all viruses follow this pattern, and it remains an assumption^41^. Additional work is required to better understand the extent to which viral material collected in respiratory samples describes the viral population in a host.

Our results provide a novel perspective upon the impact of chronic infection upon the global SARS-CoV-2 viral population. Non-trivial population structure was associated with at least one subpopulation having an inferred rate of evolution substantially greater than that of the global population. Our detection of rapidly-evolving subpopulations within chronically infected hosts highlights the potential contribution of chronic cases to the evolution of the global SARS-CoV-2 population. The increased rates of evolution inferred by our method, arising from the account made for population structure, suggest that this potential may previously have been underestimated. Our study was unable to evaluate which of the variants generated during the course of infection, if any, were transmitted to other individuals. Further work is required to combine methods for studying within-host rates of evolution with larger and more descriptive datasets, to better understand the role of vaccination and antiviral therapies upon within-host evolution, and more broadly to understand existence and consequences of virus compartmentalisation in immunocompromised hosts.

## Methods

### Sequence data

Our data comprised samples collected from nine patients who were treated for SARS-CoV-2 infection at Addenbrookes Hospital in Cambridge, UK. Identified positive samples were collected as part of a broader effort to investigate suspected hospital-acquired cases of infection^42^. In investigating suspected cases a multiplex PCR-based approach was used according to the modified ARTIC version 2 protocol with version 3 primer set, and amplicon libraries sequenced using MinION flow cells version 9.4.1 (Oxford Nanopore Technologies, Oxford, UK). Patients were selected from a dataset of cases of infection with samples collected prior to the end of October 2020, for whom i) at least four high-quality genome sequences were available and ii) the time interval between the collection of samples for the first and the last sequences was at least 21 days. Date of symptom onset was derived from the ISARIC study^43^ for eight patients. Consensus level sequence data generated via nanopore sequencing were available for all nine patients^42^. To supplement these data we reanalysed short-read sequence data describing in more detail one of the nine cases of infection; a previous analysis of these data was published elsewhere^10^. Samples were collected via swab, sputum sample, or via aspirate. Diagnostics were carried out either using qPCR or using the Hologic Panther Fusion approach^44^. CT values for patients, where these were available, are shown in Supplementary Figure 7.

### Phylogenetic trees

Phylogenetic trees were created with iQTree2^45^ and were visualised using iToL^46^.

### Mapping consensus viral sequences

We developed the Blanche software package to perform sequence-based cartography, inspired by the use of similar methods for plotting genetic and antigenic data^47,48^. Blanche carries out a simple dimension reduction calculation to facilitate the plotting of viral genome sequence data collected from individual patients. Given a set of viral sequences we first calculated a matrix *D^H^* of Hamming distances between sequences. Ambiguous nucleotides in the data were represented in our calculation by their expected value. Where a sequence contained an ambiguous nucleotide (e.g. N) at a variant site in the population, this nucleotide was set to the expected value given the composition of other sequences from the same individual. For example, if at a site there were three unambiguous observations of an allele A, and two observations of an allele C, an ambiguous nucleotide would be defined as (0.6 A, 0.4C). The distance from this nucleotide to an A would then be equal to 0.4, while the distance from this nucleotide to a C would be 0.6.

Given values for the matrix *D^H^*, we then used a simple optimisation process to identify a set of points in two-dimensional Euclidean space, with matrix of Euclidean distances between points *D^E^*, so as to minimise the sum of the squared distances between matrix elements

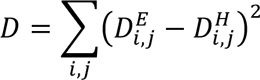

In this calculation two-dimensional Euclidean space was chosen to provide a simple representation of points on a page or screen. We note that for many datasets the minimum value of *D* was non-zero: Our representations provide an approximate picture of the relationships between genome sequences.

### Inference of subpopulations and rates of evolution

We inferred parameters for models of viral populations which successively included higher numbers of distinct subpopulations, using model selection via the Bayesian Information Criterion^49^ to identify the best fit to the data.

Common to each model we identified variant positions in the consensus viral sequences from each time point and converted these to binary code, where a 0 represented the first observed allele at a site in the genome and a 1 represented the variant allele. Ambiguous nucleotides were marked as an N, while non-variant sites were removed from consideration. This process resulted in a binary matrix of sequence data, with columns representing variants and rows representing individual sequences.

To the data we added an initial consensus sequence at time zero. By default this initial consensus was equal to the sequence of the first sample observed within the patient. However, changes to this default were accounted for in a model-specific way, described below.

Samples were then divided into subpopulations; our models allowed for between one and four such populations. Each population shared the initial consensus sequence, which described the state of the system at time zero. Constraints were applied in the division of samples, requiring that at least one subpopulation had to be represented by at least three distinct samples, and that, where the sequences of two samples shared at least two variants relative to the consensus, these samples had to be within the same subpopulation; this latter constraint acts like a weak form of parsimony.

Having defined subpopulations, variant positions were re-calculated, specific to each subpopulation. Variant positions were then split into what we termed ‘fixation’ and ‘fluctuation’ sites. For each column in our matrix of sequence data, now defined for a specific population, we calculated the Hamming distance between the column vector and the elements of two sets of vectors of equivalent length. Where there were *k* sequences in our subpopulation, the set *V_0_* comprised two vectors of length *k*, one consisting of all zeros, and the other consisting of all ones. The set *V_1_* comprised *k*-1 vectors, with vector *l* in the set containing *l* zeros followed by *k*-*l* ones, representing a fixation event occurring between sample *l* and sample *l*+1. Where the minimum Hamming distance between a column and a vector in set *V_1_* was smaller than the minimum Hamming distance between a column and a vector in set *V_0_*, a variant position was classified as a fixation; otherwise the position was classified as a fluctuation.

For each fixation we next identified the time interval in which a fixation took place as that between the first observation of a 1 in at a variant site, and the previous observation from that subpopulation. For any given time interval between samples, we could thus calculate the number of fixations occurring in that interval. The number of fluctuations in a sample was defined as the number of observations of a 1 in fluctuation positions, plus the number of observations of a 0 at positions in fixation positions after the time of a fixation event.

Given these data, we calculated the likelihood of a specific rate of evolution. We specifed a rate of evolution λ, measured in units of number of fixations per day, and an error parameter ε, measured in units of the number of errors per sequence. The likelihood of these two rates was then specified by

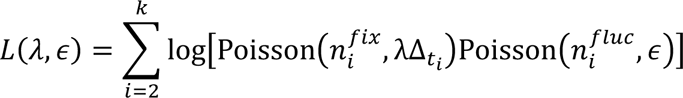

Where 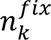 is the number of fixations in the time period between the collection of sample *i* and the previous time-point, Δ_*ti*_ is the time in days between the collection of sample *i* and the previous time-point, 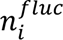 is the number of fluctuations in sample *i*, and

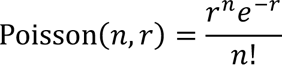

Fluctuations in the first time point were not considered, as this was defined by the initial consensus.

The log likelihood of a model was calculated as the sum of the log likelihoods *L(*λ,ε) across all subpopulations. An improvement in the Bayesian Information Criterion of 10 units was required for the acceptance of a more complex model, representing a conservative cutoff for the acceptance of additional subpopulations. In our model we inferred independent values of λ for each subpopulation. The value of ε was fixed to a constant value of 0.207 nucleotide errors per sequence, using a value inferred from a large set of sequences from 136 patients at Addenbrookes Hospital, sequenced using the same protocol^18^. That dataset imposed requirements on the completeness of a viral sequence; investigation of those data did not identify a strong effect of CT value upon the number of nucleotide errors in a sequence^18^.

### Stochasticity in defining fixation events

While the emergence of a novel mutation in the last sample to be collected from a population was defined by the method above as a fixation event, variants arising in this manner are formally indistinguishable from changes occurring through sequencing error. These events were treated in a stochastic manner. Our likelihood calculation deals only with the total number of fixations or fluctuations, rather than their location. If for a population there were 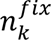 fixations in the final time interval, we considered the ensemble of cases for which there were s fixations, and 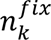-*s* additional fluctuations observed in the final time point. The maximum likelihood model was defined as the maximum likelihood calculated for any case within the ensemble.

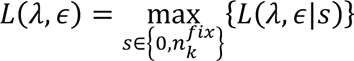

### Defining the initial consensus

Our approach assumed that any subpopulations within a host were founded from a common initial consensus sequence, rather than by viruses with distinct sequences. The initial consensus sequence was defined in a model-specific way, as follows:

### Model I: One subpopulation

Where data were available, the time of symptom onset for a patient was denoted as day zero; if not then the time of the first collected sample was denoted as day zero. If the first available sample was collected on day zero of infection, this sample was used as the initial consensus; evolution was modelled from this point forwards. In the event that the first sample was not collected on day zero, the potential exists for substitutions to have occurred in the viral population prior to the collection of the first sample. We denote by *n_pre_* the number of substitutions prior to the collection of the first sample, and optimised over this parameter.

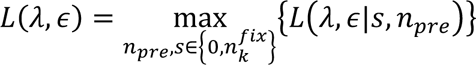

### Model V: Two subpopulations

Model V describes the evolution of two distinct viral populations, which are separated at the moment of initiating infection. Within this model we have two subpopulations, each of which has a first-collected sequence; these two sequences may differ by some number of variants. An ensemble of models were generated in which the consensus sequence was one of a set of possible sequences including the two first-observed sequences and any possible sequence intermediate to the two.

### Model Y0: Three subpopulations

Model Y0 describes the evolution of three populations with a common initial consensus. Given an infinite sites assumption, that variants may be gained but never lost, the initial consensus sequence in this model could not contain any of the variants distinct to the initial populations, so was well-defined.

### Model X0: Four subpopulations

Model X0 describes the evolution of four populations with a common initial consensus. As with model Y0, the initial sequence was well-defined.

### Parameter calculations

Maximum likelihood values of rate parameters were calculated using a simple optimisation process for any given assignment of samples to subpopulations. All possible assignments of samples to subpopulations were considered, in a systematic search process. Upper (and lower) bounds for single rate parameters were conducted using a ratcheting process whereby a parameter was fixed to only increase (or decrease) while other parameters were free to change, subject to the likelihood being within two units of the maximum likelihood. Where, given the maximum likelihood assignment of samples to subpopulations, different initial sequences or values of the parameter *s* led to likelihoods within two units of the maximum likelihood, calculations were repeated across all such initial sequences and values, finding extreme values subject to the likelihood constraint. The identification of plausible combinations of rate parameters, used to generate Figure 2B, was conducted on a grid, altering parameters by units of 0.002, and finding parameter combinations that produced likelihoods within 2 units of the maximum likelihood; as previously this approach explored all plausible initial sequences and values of *s*.

The probability of a specific variant fixing in a population was calculated across the ensemble of all potential model outcomes. Different assignments of samples to populations, different starting sequences, and different decisions about fixations in the last time point can all affect whether a fixation event was inferred to exist. The probability of a specific variant fixing was calculated as the weighted sum over all model outcomes of a binary parameter denoting whether the variant fixed, with weightings given by the likelihood function.

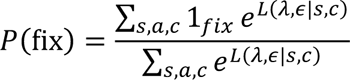

Where *c* is the initial consensus sequence, and *1_fix_* equalled one if the variant fixed given s and *c*, and zero otherwise.

Rates of substitutions at synonymous and non-synonymous sites were calculated in terms of the number of fixation events at each type of site per day of evolution. Suppose that P(fix)*_i,j,k_* denotes the probability that variant *k* fixes in subpopulation *j* in patient *i*. Then the rate of non-synonymous evolution for this population was calculated, in units of substitutions per nucleotide per day, as

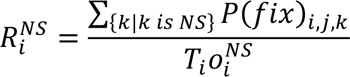

Where o denotes the number of sites in the genome that would provide non-synonymous mutations, and *T_i_* is the period of time modelled within individual *i*. The rate of evolution at synonymous sites was calculated in a similar manner.

### Simulated data

We generated simulations describing the accumulation of mutations in simple and mixed populations. Simulations were analysed using IVY, making use of the GNU parallel software package^50^. In a simple population, evolution was modelled as occurring via a simple process of the gain of mutations. The initial sequence of the population was specified as the Wuhan Hu-1 strain of the SARS-CoV-2 virus. The population was then modelled in terms of a consensus sequence.

Sequences were sampled from our population at regular times, spaced at intervals Δ*t*. At each sampling point, the viral sequence gained a number of consensus mutations defined as the outcome of a Poisson process with rate λ. Specifically, the number of changes to the sequence was calculated as

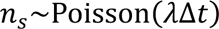

Changes in the sequence were modelled to occur at uniformly randomly sampled positions within the sequence. No distinction was made between changes that caused synonymous, non-synonymous, or nonsense mutations.

Further to the changes in the viral population, we modelled errors as arising in the sequence at rate μ, independent of time, so that the observed sequence contained *n_e_* errors, where

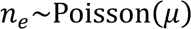

Errors also occurred at uniform random positions in the sequence.

By default we collected samples at seven time points, with the first sample collected at time zero, and Δ*t* = 5 days. Simulations were calculated for values of λ in {0.1, 0.2, 0.3} day^-1^ and with μ=0.206845

Samples were collected at seven five-day intervals, beginning at day zero, with the final sample collected on day 30. Simulations were calculated for values of λ in {0.1, 0.2, 0.3} per day and μ=0.207.

Mixed populations were generated in a similar manner to simple populations, but with two or three underlying viral populations that shared an initial consensus sequence. Each population evolved independently of the other, with an independent rate of evolution. At each sampling point the consensus sequence of one of the underlying populations was recorded, subject to error. The schedule of which population a consensus was collected from at each sampling point was randomly chosen to have an equal probability of sampling either population at a given time point, but with the restriction that at least two samples were collected from each population across the set of samples collected. Samples from simulations involving three underlying populations were collected at ten five-day intervals, with the final sample collected on day 45.

### Availability of code

Blanche can be downloaded from the repository https://github.com/cjri/Blanche. IVY can be downloaded from the repository https://github.com/cjri/IVY.

### Ethics statement

This study was conducted as part of surveillance for COVID-19 infections under the auspices of Section 251 of the NHS Act 2006. The ISARIC/WHO Clinical Characterisation Protocol UK (ISARIC CCP-UK) was approved by the Oxford C Research Ethics Committee (reference: 13/SC/0149) and by the Scotland A Research Ethics Committee (reference 20/SS/0028). The COG-UK study protocol was approved by the Public Health England Research Ethics Governance Group (reference: R&D NR0195).

## Acknowledgements

This work was supported by funding from the UK Medical Research Council (MC_UU_00034/6, 207498/Z/17/Z and 206298/Z/17/Z). We thank Iain Barrass for research computing support. This work uses Data / Material provided by patients and collected by the NHS as part of their care and support #DataSavesLives. The Data / Material used for this research were obtained from ISARIC4C. The COVID-19 Clinical Information Network (CO-CIN) data was collated by ISARIC4C Investigators. Data and Material provision was supported by grants from: the National Institute for Health Research (NIHR; award CO-CIN-01), the Medical Research Council (MRC; grant MC_PC_19059), and by the NIHR Health Protection Research Unit (HPRU) in Emerging and Zoonotic Infections at University of Liverpool in partnership with Public Health England (PHE), (award 200907), NIHR HPRU in Respiratory Infections at Imperial College London with PHE (award 200927), Liverpool Experimental Cancer Medicine Centre (grant C18616/A25153), NIHR Biomedical Research Centre at Imperial College London (award IS-BRC-1215-20013), and NIHR Clinical Research Network providing infrastructure support. The views expressed in this report are those of the authors and not necessarily those of ISARIC4C, NIHR, MRC, or PHE. ISARIC4C welcomes applications for data and material access through our Independent Data and Material Access Committee (https://isaric4c.net).

## Supplementary Figures

**Supplementary Figure 1:**
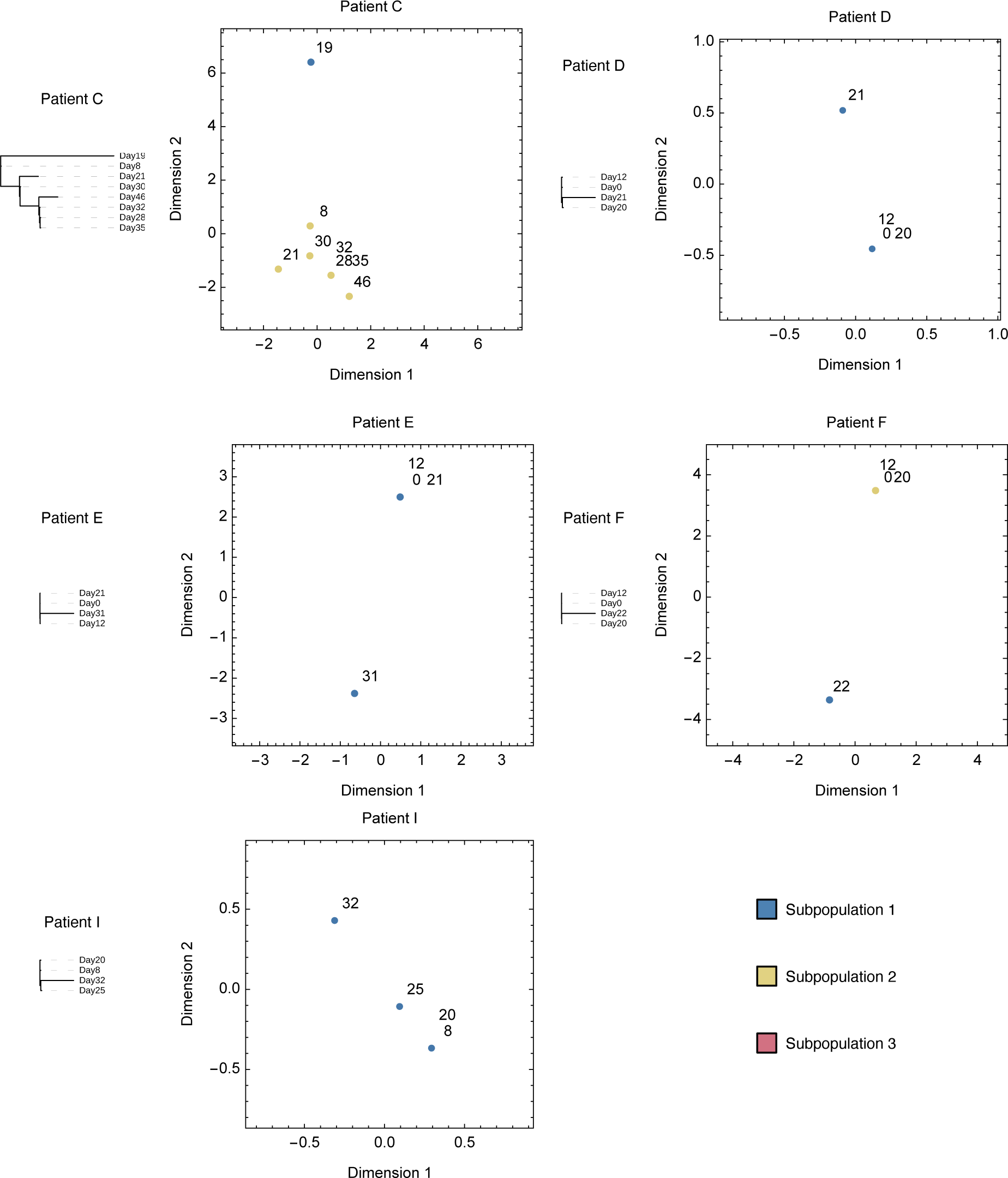
Phylogenetic trees and sequence maps for data from other patients in the study.

**Supplementary Figure 2:**
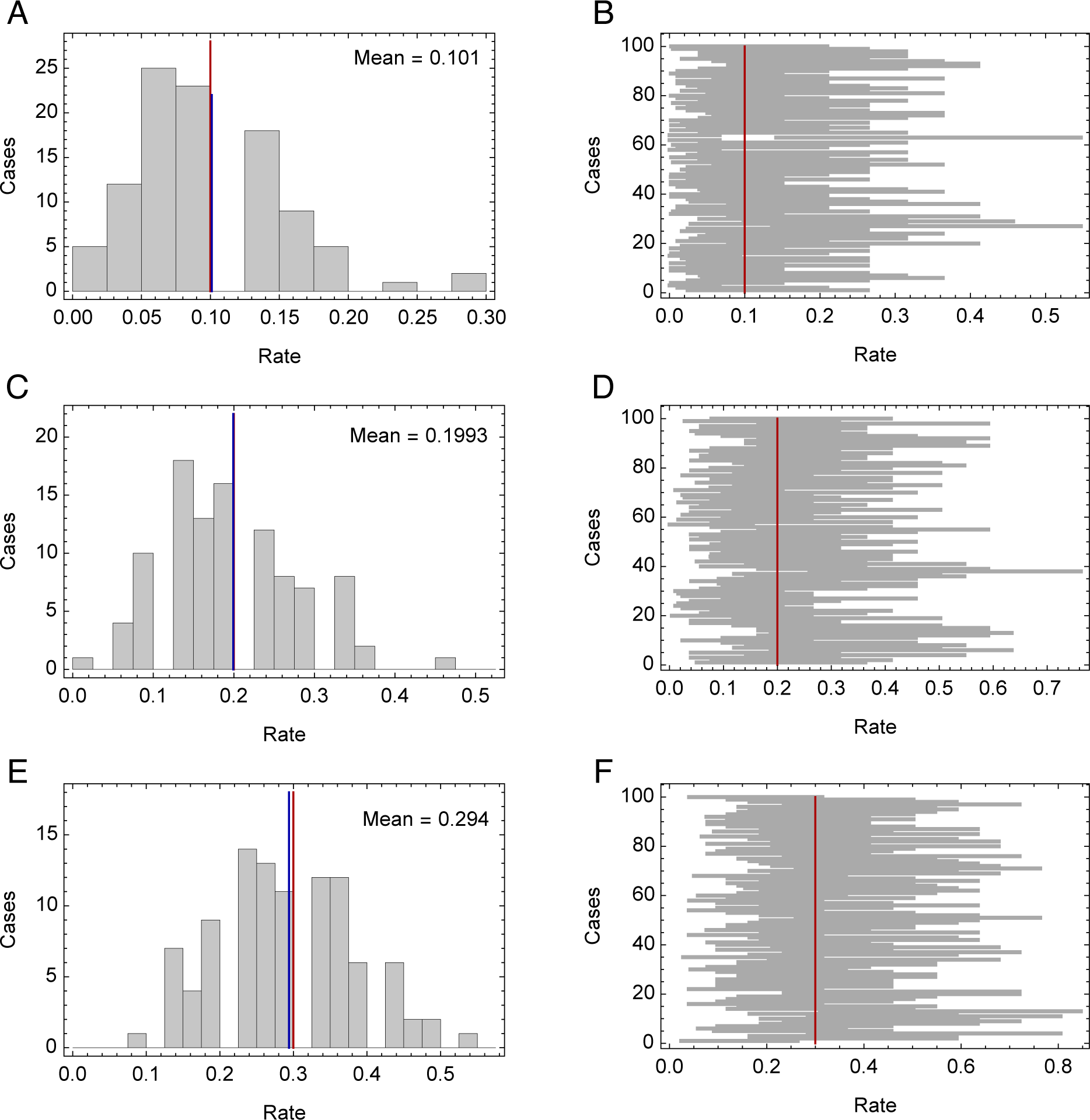
Inferred rates of evolution from simulated data. Simulations described simple populations with an underlying rate of evolution, simulated for 30 days. The actual number of mutations gained by a population during this period is Poisson distributed according to the product of the rate and time. **A.** Inferred rates of evolution (gray bars) given a rate of 0.1 per day. The mean inferred rate across 100 populations (vertical blue line) is very similar to the actual rate (vertical red line). **B.** Inferred 95% confidence intervals for these inferences. The correct rate of evolution (vertical red line) was contained within 93 of 100 intervals. **C.** Inferred rates of evolution given a rate of 0.2 per day. The mean inferred rate (vertical blue line) is very similar to the actual rate (vertical red line). **D.** Inferred 95% confidence intervals for these inferences. The correct rate of evolution (vertical red line) was contained within 98 of 100 intervals. **E.** Inferred rates of evolution given a rate of 0.3 per day (gray bars). The mean inferred rate (vertical blue line) is very similar to the actual rate (vertical red line). **F.** Inferred 95% confidence intervals for these inferences. The correct rate of evolution (vertical red line) was contained within 98 of 100 intervals.

**Supplementary Figure 3:**
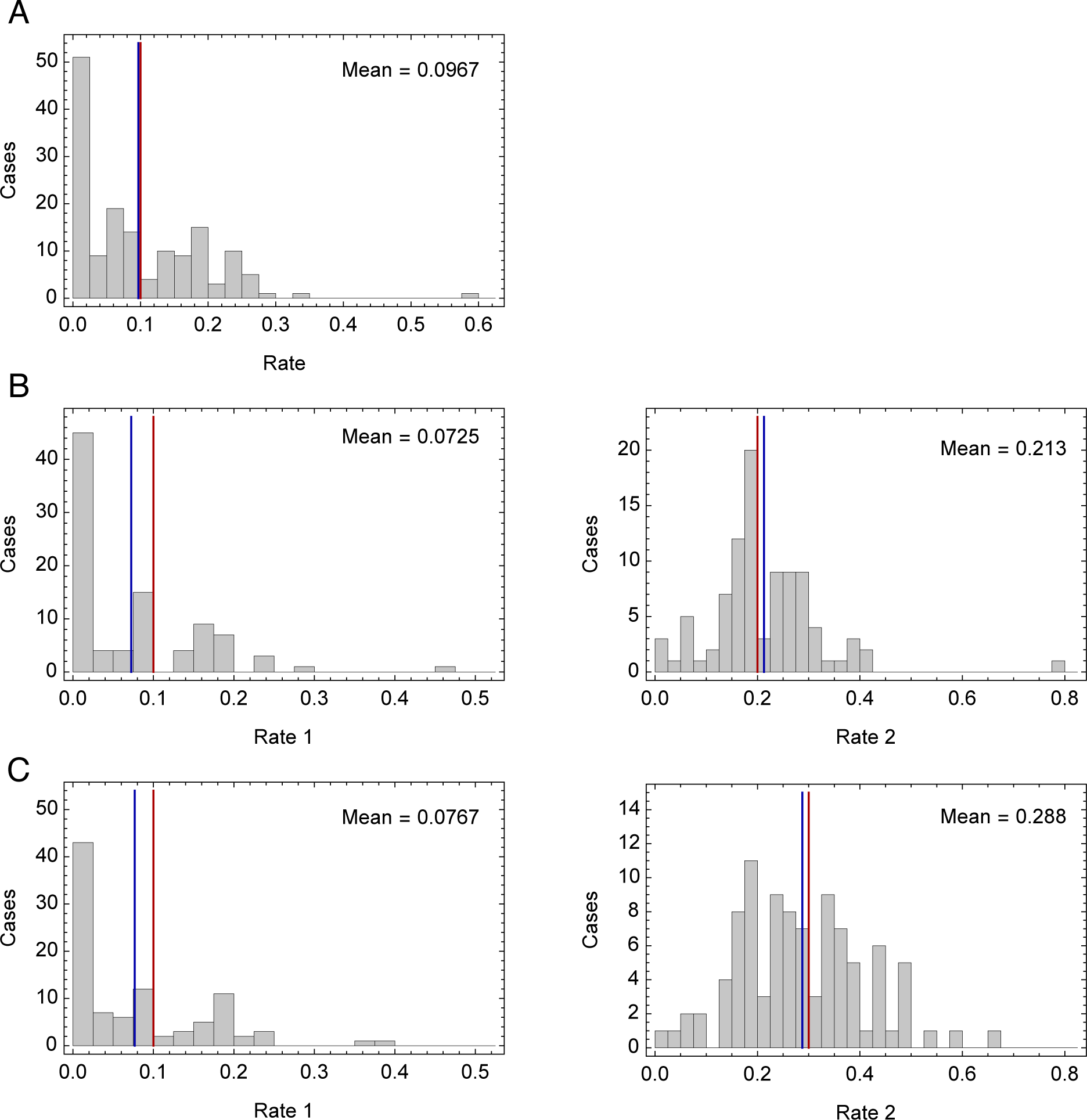
Inferred rates of evolution from simulated data. Data describe populations with two subpopulations, simulated for 30 days. **A.** Inferred rates of evolution (gray bars) where viral populations evolve at rate 0.1. The mean inferred rate (vertical blue line) and actual rate (vertical red line) are shown. **B.** Inferred rates of evolution (gray bars) where viral populations evolve at rates 0.1 and 0.2. The mean inferred rate (vertical blue line) and actual rate (vertical red line) for each population are shown. **C.** Inferred rates of evolution (gray bars) where viral populations evolve at rates 0.1 and 0.3. The mean inferred rate (vertical blue line) and actual rate (vertical red line) for each population are shown.

**Supplementary Figure 4:**
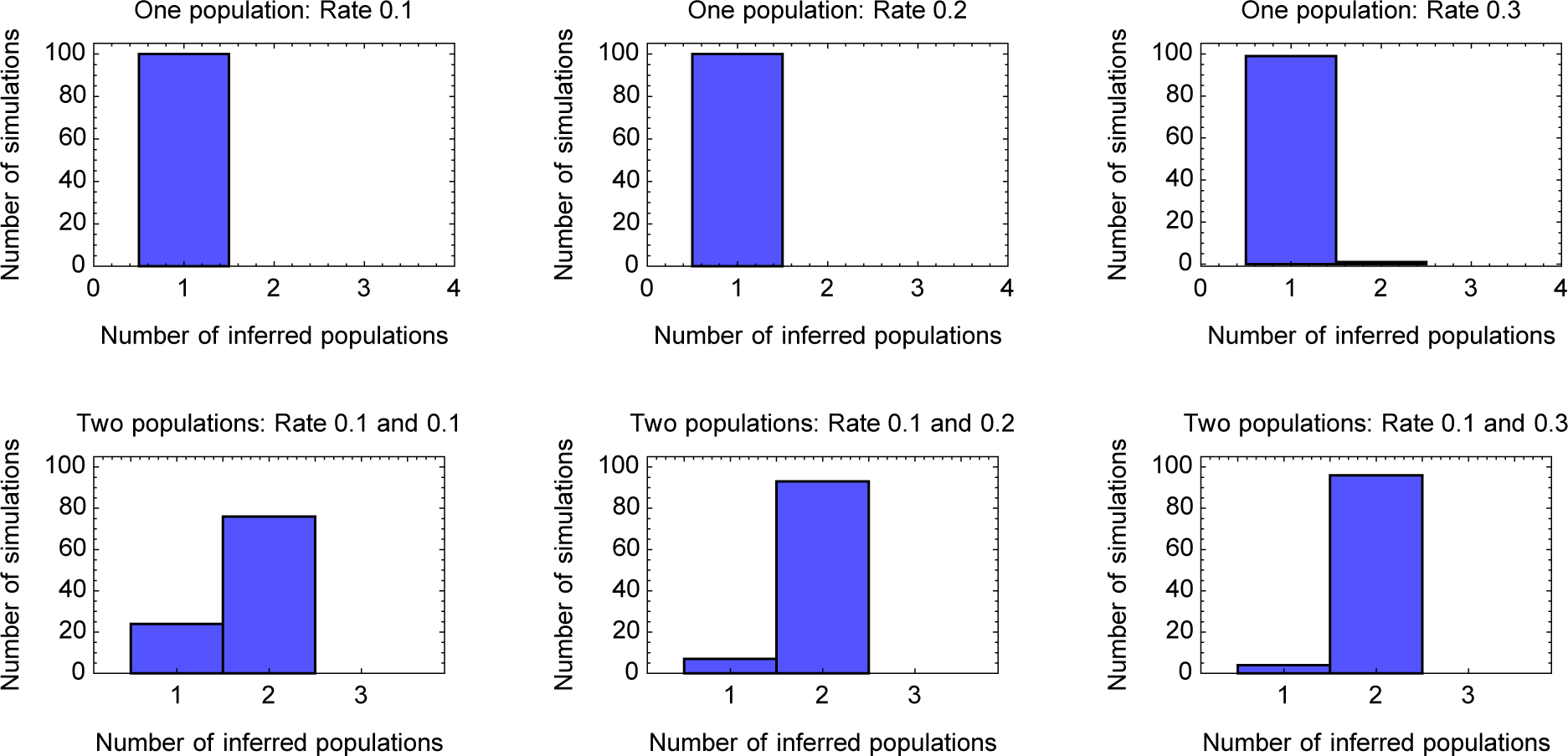
Numbers of populations inferred from simulated data. Data describe populations with one or two subpopulations, simulated for 30 days. Under-calling of distinct populations occurs in up to 25% of inferences for the case where the rates of evolution of the populations are identical, but over-calling of distinct populations was rare.

**Supplementary Figure 5:**
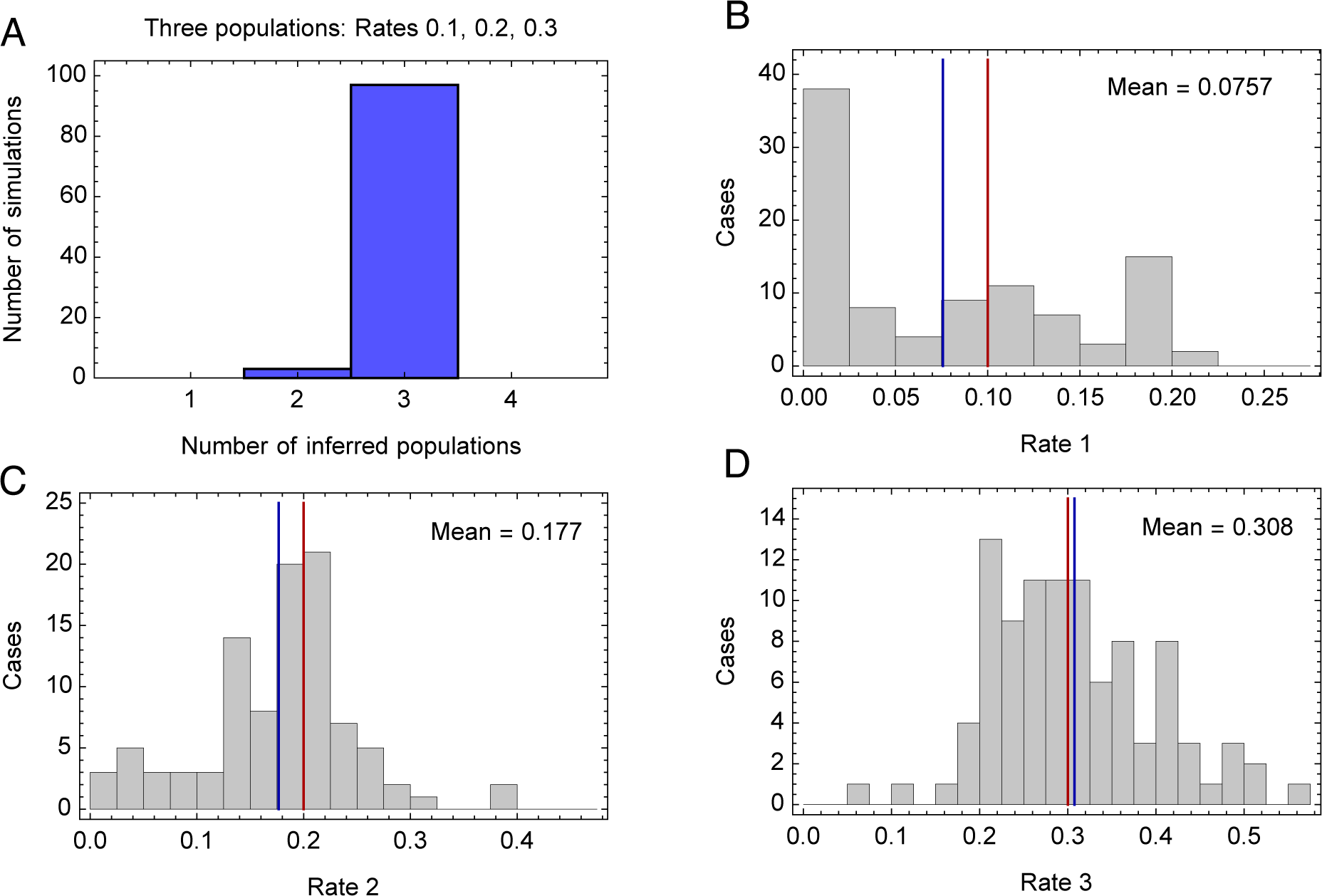
Statistics from applications to simulated data with three subpopulations. A. Number of subpopulations identified by the model. Three subpopulations were identified in 97 out of 100 simulations. **B.** Smallest fitness parameter inferred by the model shown as a histogram (gray bars). The red line and blue line show the simulated rate and the mean inferred rate of evolution respectively. **C.** Intermediate fitness parameter inferred by the model shown as a histogram (gray bars). The red line and blue line show the simulated rate and the mean inferred rate of evolution respectively. **D.** Largest fitness parameter inferred by the model shown as a histogram (gray bars). The red line and blue line show the simulated rate and the mean inferred rate of evolution.

**Supplementary Figure 6:**
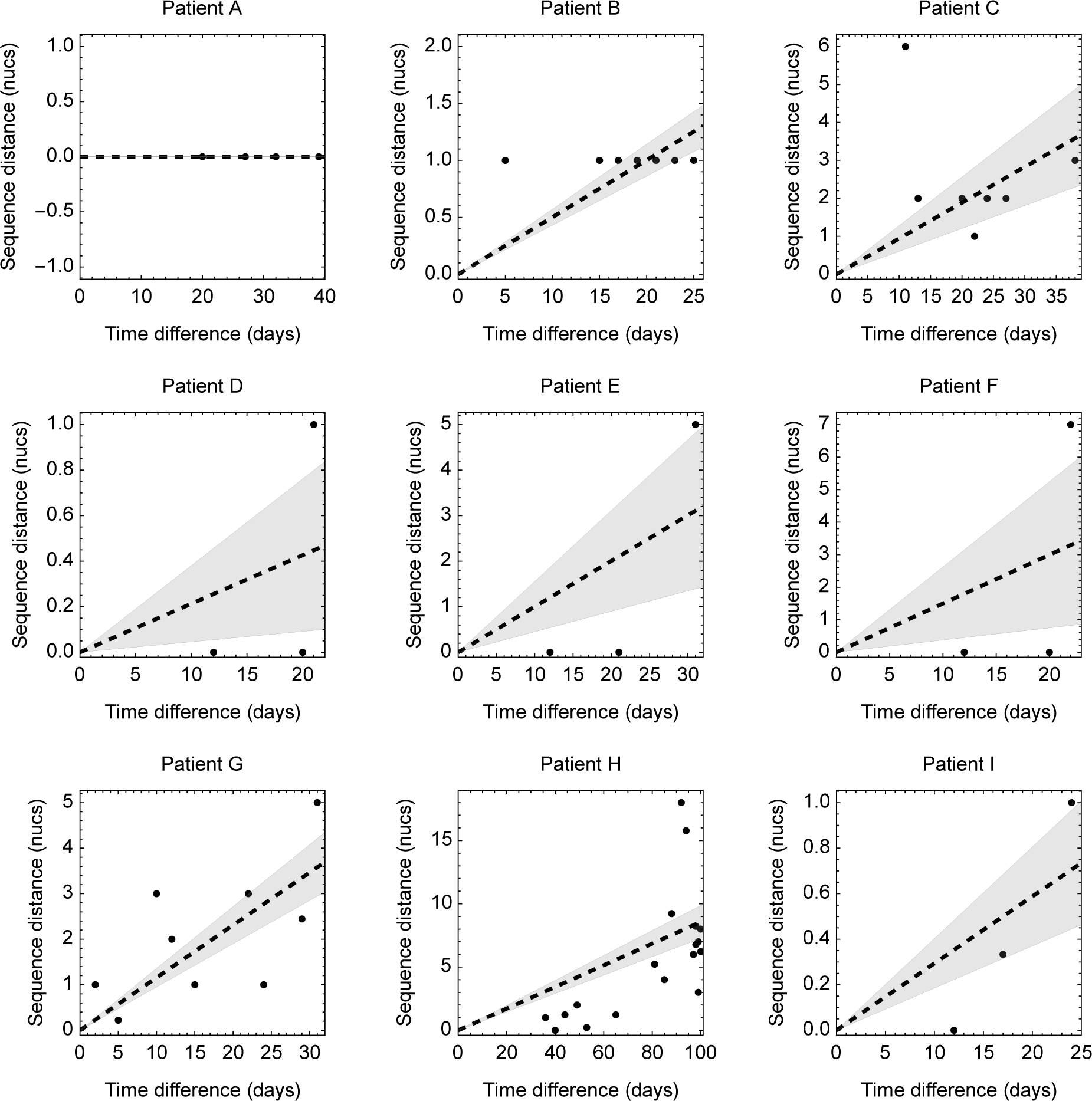
Rates of evolution inferred using a simple method of linear regression. Individual sequences are shown as dots. Non-integer sequence distances were sometimes identified in the case of missing nucleotide data. The black dashed line for each case shows a linear regression model fitted to the data. The gray shaded region shows confidence intervals for this model.

**Supplementary Figure 7:**
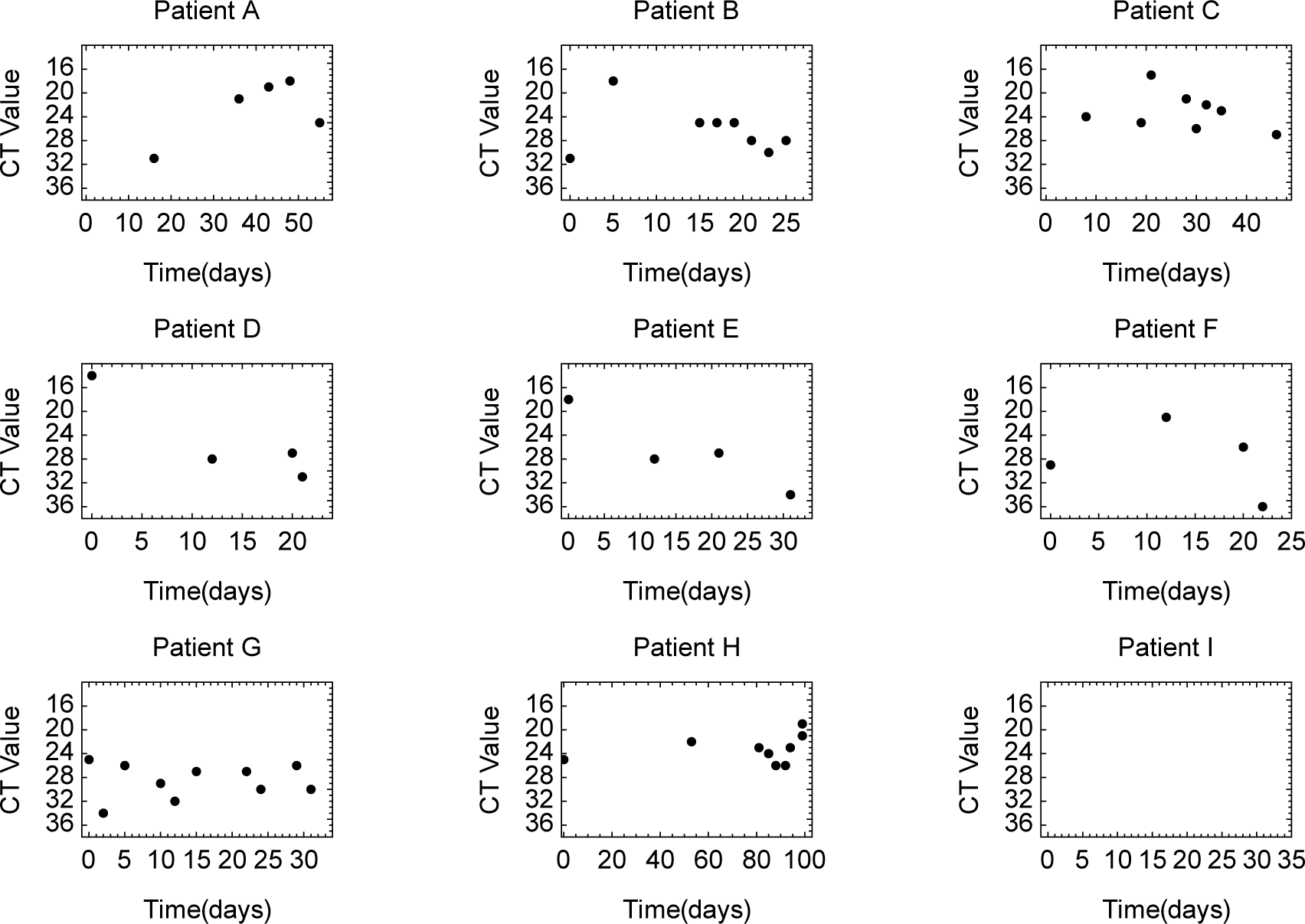
CT values for samples collected from patients, where recorded. Times are shown relative to the time of symptom onset, where known, or to the time of the first collected sample. Samples from Patient I were generated using the Panther platform, such that no CT values were recorded.

## Supplementary Tables

**Supplementary Table 1:** Counts of swabs and sputum samples from patient G, alongside the subpopulations to which they were assigned. No significant association was identified between subpopulation and type of sample (p-value 0.21, Fisher exact test). Samples of unknown sampling method (1 sample) were neglected from the analysis.

|  | Swab | Sputum |
| --- | --- | --- |
| Subpopulation 1 | 2 | 1 |
| Subpopulation 2 | 1 | 5 |

**Supplementary Table 2:** Counts of swabs and sputum samples from patient H, alongside the subpopulations to which they were assigned. No significant association was identified between subpopulation and type of sample (p-value 0.35, Fisher exact test). Aspirate samples (1 sample) were neglected from the analysis.

|  | Swab | Sputum |
| --- | --- | --- |
| Subpopulation 2 | 3 | 1 |
| Subpopulation 1+3 | 14 | 1 |

**Supplementary Table 3:**
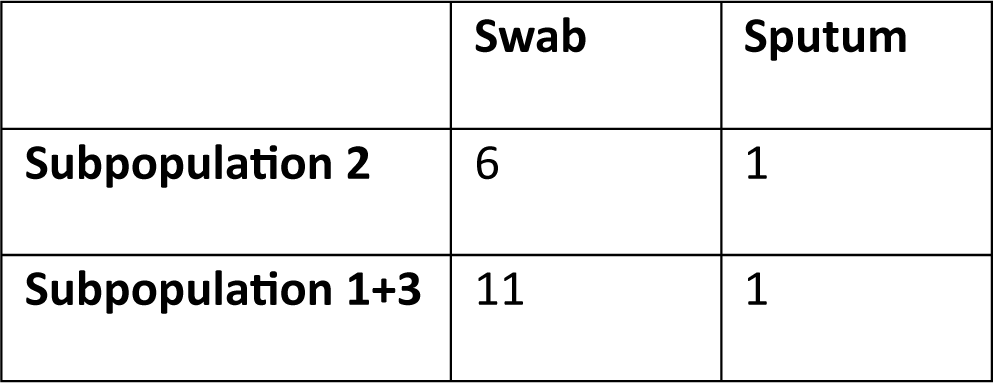
Counts of swabs and sputum samples from patient H, alongside the subpopulations to which they were assigned. No significant association was identified between subpopulation and type of sample (p-value 0.49, Fisher exact test). Aspirate samples (1 sample) were neglected from the analysis.

## Notes

### Competing Interest Statement

The authors have declared no competing interest.

